# Peptide sequencing in an electrolytic cell with two nanopores in tandem and exopeptidase

**DOI:** 10.1101/015297

**Authors:** G. Sampath

## Abstract

A nanopore-based approach to peptide sequencing without labels or immobilization is considered. It is based on a tandem cell (*RSC Adv.,* 2015, **5**, 167-171) with the structure [*cis*1, upstream pore (UNP), *trans*1/*cis*2, downstream pore (DNP), *trans*2]. An amino or carboxyl exopeptidase attached to the downstream side of UNP cleaves successive leading residues in a peptide threading from *cis*1 through UNP. A cleaved residue translocates to and through DNP where it is identified. A Fokker-Planck model is used to compute translocation statistics for each amino acid type. Multiple discriminators, including a variant of the current blockade level and translocation times through *trans*1/*cis*2 and DNP, identify a residue. Calculations show the 20 amino acids to be grouped by charge (+, −, neutral) and ordered within each group (which makes error correction easier). The minimum cleaving interval required of the exopeptidase, the sample size (number of copies of the peptide to sequence or runs with one copy) to identify a residue with a given confidence level, and confidence levels for a given sample size are calculated. The results suggest that if the exopeptidase cleaves each and every residue and does so in a reasonable time, peptide sequencing with acceptable (and correctable) errors may be feasible. If validated experimentally the proposed device could be an alternative to mass spectrometry and gel electrophoresis. Implementation-related issues are discussed.

## 1. Introduction

In nanopore sequencing, an analyte (usually a polymer) translocates through a biological nanopore embedded in a bilipid membrane (or a hole drilled through a synthetic one) separating the *cis* and *trans* chambers in an electrolytic cell with an aqueous solution of KCl and a potential difference between the two chambers. The resulting ionic current blockade is used to identify the analyte (or its components) as it passes through the pore. Instead of a pore in a membrane a graphene sheet or layer of molybdenum sulphide containing a nano-sized hole may also be used, with a transverse current passing through the analyte and a pair of transverse electrodes being used to identify monomers. In ‘strand sequencing’ [1] of DNA the analyte is a charged DNA molecule, in ‘exosequencing’ [2] it is charged bases (actually mononucleotides) cleaved by an exonuclease adjacent to the pore in *cis*. Nanopore-based sequencing of single- or double-stranded DNA (ss- or ds-DNA) has been studied extensively (see review [3]), with one implementation in beta-test mode [4]. In contrast, sequencing of proteins or peptide strands using nanopores is still in an early stage [5], in part because the problems it faces are much more severe than those in DNA sequencing.

This report looks at the possibility of a nanopore-based method to sequence a peptide. It is centered on a modified version of a tandem electrolytic cell previously proposed for DNA sequencing and modeled mathematically [6]. That model is extended to the modified tandem cell, whose analysis suggests that peptide sequencing with a nanopore may be feasible, provided the exopeptidase functions as required by the proposed method. The original version has two pores in tandem with an exonuclease attached to the downstream side of the first pore. The enzyme is designed to cleave the leading base from a single strand of DNA that is drawn into and through the first pore by a potential difference across the cell. The cleaved base translocates to and through the second pore and is detected based on the current blockade it causes there. The cell considered here is similar, with an exopeptidase in place of the exonuclease to successively cleave leading amino acids (or residues, the two terms are used interchangeably below) in the amino acid chain. Sequence identification is based on the use of multiple discriminators, including a variant of the current blockade level and the translocation times of a cleaved residue through *trans*1/*cis*2 and DNP. If experimentally validated this approach could lead to an alternative to mass spectrometry (ESI/MALDI) [7] and gel electrophoresis [8].

The following summarizes the content of this report. Section 2 presents a brief review of nanopore-related studies of proteins/peptides and the potential use of nanopores in protein/peptide sequencing. Section 3 describes a tandem cell with exopeptidase for sequencing a peptide without any labels or immobilization. Section 4 presents results from a mathematical model based on [6]. Section 5 presents an analysis of the model and discusses conditions for effective peptide sequencing with a tandem cell. Among other things, it examines the use of multiple discriminators to enable identification of more monomer types than previously considered, as well as necessary conditions for residues to be sequenced in the correct order. Section 6 looks at a range of implementation issues. An Appendix contains tables of calculated data.

## 2. Nanopores for peptide sequencing

Most protein sequencing (more correctly peptide sequencing) is currently based on peptide ionization and mass spectrometry (ESI/MALDI) [7] or gel electrophoresis [8]. In recent years there has been an increasing number of investigations of nanopores for peptide identification and analysis. Most of this work does not involve residue-level sequencing but is concerned with other aspects such as protein unfolding [9], identification of whole proteins [10] or domains within [11], detecting modifications such as phosphorylation [12], or conformation studies [13]. A recent report [14] describes the use of transverse electrodes and residue-specific detector molecules attached to the electrodes to measure a transverse tunneling current through a single amino acid in the peptide as it translocates through the pore. The current record is then used with a machine learning algorithm to identify the sequence of amino acids crossing the junction.

In general any attempt to sequence proteins using nanopores has to consider the following: 1) unlike DNA, which carries a negative electric charge in its backbone, only 5 of the 20 individual amino acids that make up proteins are charged (2 are negative, 3 positive), the other 15 being neutral [8]; therefore depending on the sequence a peptide may carry only a small effective charge, which may not be sufficient to move or enable easy detection in an electric field; 2) proteins in their native state have secondary and tertiary folds and therefore need to be unfolded before sequencing can begin (ds-DNA has a similar unzipping problem, which can be resolved with a nanopore [15]); 3) runs of identical residues (homopolymers) are not easily resolved (this is also a problem for strand sequencing of DNA); 4) proteins are not easily replicated; in comparison DNA can be reproduced in large amounts using the polymerase chain reaction (PCR) [8]; 5) if the sequencing is based on cleaving of a strand the original sequence is not easily reconstructed, whereas in exonuclease sequencing of ss-DNA [2,6] re-sequencing of the DNA strand from the individual cleaved nucleotides can be done with a template and an enzyme motor attached to a nanopore [16]; and 6) attempts to use methods similar to DNA sequence extraction methods in which Markov-Viterbi models or neural-net-based algorithms [17,18] are used to identify bases from the current signal due to a segment of k bases (k-mer) rather than a single base have to contend with the much larger number of amino acid types (20 versus 4 in DNA); thus with k=2 the number of blockade levels to distinguish is 400 (compared with 16 in DNA), and 160000 with k=4 (compared with 256 in DNA).

Some of the above problems can be alleviated: 1) neutral molecules can be made mobile in an electric field if a hydraulic pressure gradient is added [19-21]; 2) folded proteins can be unfolded using a nanopore [9]; 3) the homopolymer problem can be solved in part by breaking up the peptide into individual residues so that the identification of each residue is, generally speaking, not influenced by its neighbors in the chain (similar to exonuclease sequencing of DNA [2] with the tandem cell [6]) because the ionic current returns to the baseline value between successive residues; and 4) multiple discriminators based on different measured data may be used to identify a larger number of monomer types. The next section describes a modified tandem cell for peptide sequencing based on some of these notions.

## 3. A tandem cell for sequencing a peptide strand

Figure 1 shows a schematic of the modified tandem cell that is based on the generic form [*cis*1, upstream pore (UNP), *trans*1/*cis*2, downstream pore (DNP), *trans*2] [6] and has a similar geometry. An exopeptidase (amino or carboxyl) is attached to the downstream side of UNP. A potential difference V_05_ (normally > 0) is applied between *cis*1 and *trans*2 over the five sections; most of it (~98%) drops across the two pores [3].

**Figure 1.**
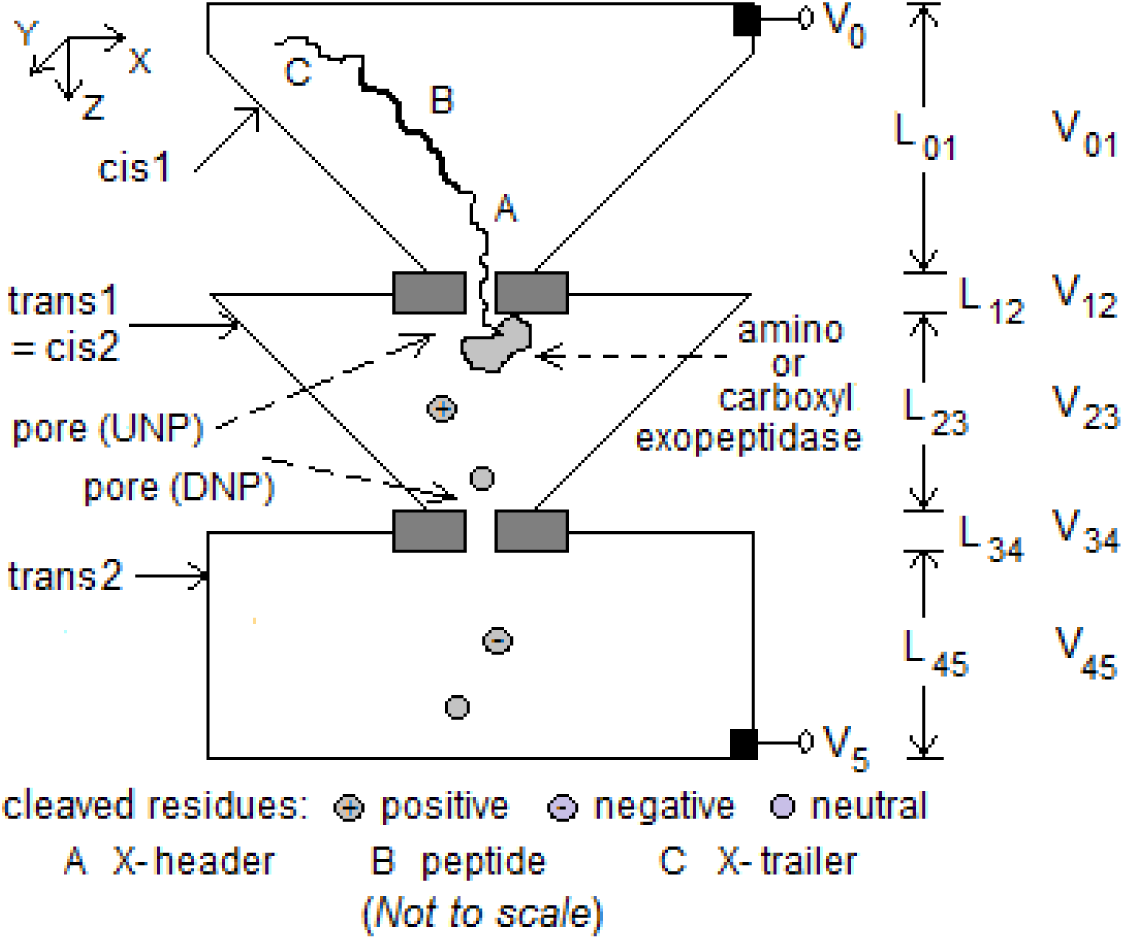
Schematic of modified tandem cell for peptide sequencing with five pipelined stages. Dimensions considered: 1) *cis*1: box of height 1 μm tapering to cross-section 100 nm^2^; 2) UNP: length 10-20 nm, diameter 10 nm; 3) *trans*1/*cis*2: box of height 1 μm tapering from 1 μm^2^ cross-section to 10 nm^2^; 4) DNP: length 10-20 nm, diameter 3 nm; 5) *trans*2: box of height 1 μm, side 1 μm. Exopeptidase covalently attached to downstream side of UNP. Electrodes at top of *cis*1 and bottom of *trans*2. V_05_ ≈ 0.4 V.

Translocation of analyte molecules in the tandem cell is primarily based on diffusion supplemented by drift due to the electric field E resulting from V. The diffusion-drift process can be modeled with a Fokker-Planck equation, and the mean and standard deviation of the times taken by the particle to translocate through a chamber (*cis* or *trans*) and through a pore calculated. With the z axis parallel to the pore axis and directed from *cis*1 to *trans*2, a negatively charged particle experiences a positive drift velocity v_z_ (= μE, where μ is the particle mobility) due to E = V/L, which reduces mean translocation time. If it is positively charged the drift is in the opposite direction, and the mean translocation time increases. The field has no effect on neutral residues.

The behavior of the proposed structure can be described as follows. A peptide with a poly-X, where X = negatively or positively charged amino acid (X− = Glu or Asp; X+ = Lys, Arg, or His), leader and trailer to induce entry into UNP (V_5_ > V_0_ or V_5_ < V_0_ respectively), is drawn into UNP when |V_05_| is sufficiently large (about 200-400 mV typically; see Figure 7 in [3]), and translocates through UNP to encounter the exopeptidase attached to the downstream side of UNP. If the exopeptidase is amino exopeptidase then the leading residues at the N-terminal of the chain are cleaved one after the other. With carboxyl exopeptidase the cleaving is at the C-terminal. (The incorrect end could enter UNP, Section 6 considers how this could be resolved.) A non-zero potential difference between *trans*1/*cis*2 and *trans*2 causes ionic current to flow through DNP. A cleaved residue passes through DNP under the influence of V_34_ and/or diffusion, causing a blockade of the ionic current. By measuring the blockade current level through DNP, the mean inter-arrival time (≈ E(T*_trans_*_1/*cis*2_)) between successive cleaved residues arriving at DNP, and the mean residence time of the cleaved residue inside DNP (≈ E(T_DNP_)) a residue can in principle be identified using these three discriminators (or their close variants). (This ignores interactions with the pore lumen and the effect of a chemical adapter [22] used for slowdown; see Section 6.)

## 4. Mathematical model

The mathematical model for the tandem cell here is very similar to that for the tandem cell proposed for exosequencing of DNA [6]. Similar to a mononucleotide in the original tandem cell, a residue is considered to be a particle that does not interact chemically with the pore lumen or the electrolyte and moves after being cleaved by the exopeptidase through a combination of diffusion and electric drift. A cleaved residue cannot regress into UNP because it is blocked by the remaining peptide in UNP. Most of the potential difference V_05_ is dropped across the two pores (V_05_ = 0.365 V, V_23_ = 1.6 mV, V_34_ = ~0.18 V). Movement of a residue, which is dominated by diffusion, can be studied via the trajectory of a particle whose propagator function G (x,y,z,t) is given by a linear Fokker-Planck (F-P) in one dimension (z) for DNP, or three (x,y,z) for *trans*1/*cis*2. The equation for G contains a drift term in the z direction that arises from the voltage difference V_05_. The drift field affects charged residues but not neutral ones. A piecewise approach is taken, with each section considered independent of the others. The behavior at the interface between two sections is examined in Section 5.4.

### 4.1 One-dimensional case

The F-P equation in the one-dimensional case can be solved in a straightforward way using methods from partial differentiation equations and Laplace transforms. Let μ be the mobility of the particle and D its diffusion constant. Following [6], the mean E(T) and variance σ^2^(T) of the translocation time T over a channel of length L that is reflective at the top and absorptive at the bottom with applied potential difference of V are given by

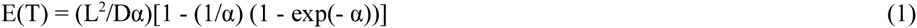

and

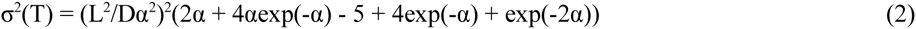

where

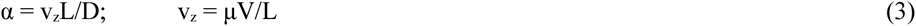

Here v_z_ is the drift velocity due to the electrophoretic force experienced by a charged particle in the z direction. For v_z_ = 0, these two statistics are

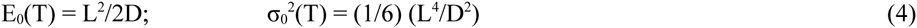

As discussed below, these formulas can be applied to all three channels: *trans*1/*cis*2 (T = T*_trans_*_1/*cis*2_; L = L_23_), DNP (T = T_DNP_; L = L_34_), and *trans2* (T = T*_trans2_*; L = L_45_). The characteristics of translocation in each channel are discussed next.

### 4.2 Characteristics of translocation of a residue

1) *Translocation of a cleaved residue through DNP*. A cleaved residue is treated as a particle that is released at the top of DNP at t = 0, reflected there at t > 0, and ‘captured’ at the bottom at t > 0. Regardless of whether a residue is charged or not the diffusion is always in the z direction because of the reflecting barrier at z = 0. (Thus a cleaved residue cannot regress into UNP because it is blocked by the remaining peptide.) With V_05_ > 0 α is positive for negative residues and negative for positive residues. The resulting mean translocation time for negative residues is reduced below that due to v_z_ = 0, and goes above for positive residues. In both cases the net translocation is in the positive z direction for the values of V_05_ in use. The electric field has no effect on neutral residues and their movement is entirely due to diffusion; therefore α = 0 for them. In summary all residues, charged or not, will move in the z direction and cause a current blockade in DNP; this, along with other measures (see Section 5.1), can be used to identify a residue. Equations 1 through 4 apply with L = L_34_.

2) *Translocation of a cleaved residue through* trans*1*/cis*2*. This is modeled in three dimensions using a rectangular box-shaped region. (The tapered geometry of Figure 1 is discussed in Section 5.4.) A particle is released at the top center of *trans*1/*cis2* at t = 0, ‘reflected’ at the top and sides of the box at t > 0, and translocates to the bottom of the compartment where it is ‘absorbed’ at some t > 0. That is, the particle is considered to be detected when it reaches z = L_23_ independent of x and y and to move into DNP without regressing into *trans*1/*cis*2. The propagator function G(x,y,z,t) can be written as the product of three independent propagator functions. It is shown in [6] that diffusion in the x and y directions has no effect on G(x,y,z,t) so that the first passage time (that is, translocation time) distribution in the three dimensional case reduces to that in the 1-d case. Thus Equations 1 through 4 apply with L = L_23_. The effect of α on charged and neutral residues is the same as in DNP.

3) *Translocation of a cleaved residue through* trans*2*. This behavior can be modeled in the same way as that of a cleaved residue in *trans*1/*cis*2.

## 5. Analysis and computational results

The ability of the tandem cell to correctly identify residues cleaved from a peptide depends on the following conditions being satisfied:

1. The tandem cell must be able to discriminate among 20 types of residues;
2. Residues must not be lost to diffusion;
3. Residues must arrive in sequence order at DNP;
4. More than one residue must not occupy DNP at any time.

Conditions 2 through 4 also serve to define the minimum interval required between successively cleaved leading residues in the peptide; this is discussed in Section 5.3.

### 5.1 Discriminating among the residue types using multiple discriminators

Most sequencing studies (see review [3]) focus on the current blockade when discriminating among monomer types. In sequencing of single strands of DNA higher-level correlations among the bases in a k-mer are extracted from the current record by complex algorithms to improve base calling [17,18]. If sequencing is based on a graphene sheet with a hole for the nanopore the discriminator used is the transverse current passing through the analyte and a pair of transverse electrodes [23]. Although the residence time of an analyte in a pore has been modeled in many studies few consider it as a discriminator, most are largely from the perspective of slowing down translocation to decrease the detection bandwidth.

By using multiple discriminators in the recorded signal it may be possible to better distinguish among monomer types and/or increase the number of types that can be identified. Thus going beyond the current blockade, analyte-specific information may also be found in the times taken for a molecule or cleaved monomer to travel to a pore and through the pore. In a tandem cell both these times (or their variants) are clearly defined (translocation through *trans*1/*cis*2 to the entrance of DNP and translocation through DNP) and can be measured. Thus three discriminators, namely the mean blockade current ratio <I/I_0_> (where I and I_0_ are the currents with and without analyte in the pore), the mean translocation time E(T*_trans_*_1/*cis*2_) from top center of *trans*1/*cis*2 to the entrance of DNP, and the mean residence time E(T_DNP_) in DNP, can in principle be used in combination for analyte identification in a tandem cell. Computation of these three discriminators is considered next.

#### a) Current blockade level inside DNP and volume excluded in a pore by an analyte particle (monomer)

Current blockade is defined by the mean blockade current ratio <I/I_0_>. For polymer sequencing based on current blockades to work there must be an ionic current (due to K^+^ and Cl^−^) between *trans*1/*cis*2 and *trans*2; thus V_34_ cannot be 0. The blockade level is influenced by many factors, one of which is volume exclusion whereby the particle reduces the pore volume available for ionic current flow. The volume exclusion ratio (VER) is defined as volume excluded/pore volume: V_excl_/V_pore_. The particle is treated as a cylinder of radius equal to the particle’s hydro-dynamic radius R_H_ [24] and height 2R_H_, the pore is a cylinder of radius r and length L. The VER is given by

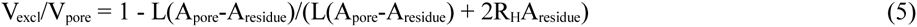

where the A’s are cross-section areas. Although it ordinarily contributes only a small fraction to the blockade ratio <I/I_0_>, the VER is used here as a placeholder and included in the discussion below for the purpose of studying the efficacy of multiple discriminators. When experimentally obtained or theoretically calculated values for <I/I_0_> become available, it may be replaced with <I/I_0_> after which the model can be revised as appropriate.

#### b) Translocation time through DNP

This is a function of the diffusion constant D_aa_ for an amino acid, its hydrodynamic radius R_H-aa_, and the drift velocity v_z_ if the residue is charged. It is also influenced by the selectivity of the pore for anions or cations (see discussion in Section 6). Additionally the translocation can be slowed down if a chemical adapter is used [22]. The mean and standard deviation of the translocation time are given by Equations 1 through 4.

#### c) *Translocation time through* trans*1*/cis*2*

The dependence on physical-chemical properties is similar to (b).

The statistics of the two translocation times for each amino acid can be calculated using Equations 1 through 4, with D and μ for an amino acid given by

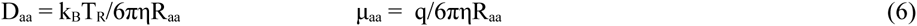

Here k_B_ is the Boltzmann constant (1.3806 × 10^−23^ J/K), T_R_ is the room temperature (298° K), η is the solvent viscosity (0.001 Pa.s), R_aa_ the hydrodynamic radius of an amino acid (usually given in angstrom (Å = 10^−10^ m)), and q is the electron charge (1.619 × 10^−19^ coulomb). Values of R_aa_ are taken from [24].

### 5.2 Computational results

Figure 2 is a scatter diagram of E(T_DNP_) vs V_excl_ / V_DNP_, while Figure 3 relates E(T_DNP_) and E(T*_trans_*_1/*cis*2_). (Calculated data can be found in Table 1 in the Appendix.) In both cases a grouping of the amino acids by electric charge (+, −, neutral) is evident, and within each group there is a monotonic ordering of the residues. The ordering property is especially useful because error correction merely requires an incorrect call to be replaced with the nearest neighbor in the ordering. (Error correction may also be enhanced by methods similar to those used in mass-spectrometry-based peptide sequencing in which pattern recognition techniques and/or correlation analysis are used with a protein sequence database to fix unknowns in a peptide fragmentation spectrum [7].) Furthermore, if the voltage V_05_ is reversed the negative and positive residues reverse position in both charts; this property can be used to advantage in sequencing as discussed below. (As an aside, a comparison with the amino acid separation spectrum obtained from ion mobility spectrometry [25], which shows a strict (mobility-based) ordering of the amino acids over drift time (with values in the milliseconds range), reveals similar tendencies between the two orderings and some overlapping segments.)

**Figure 2.**
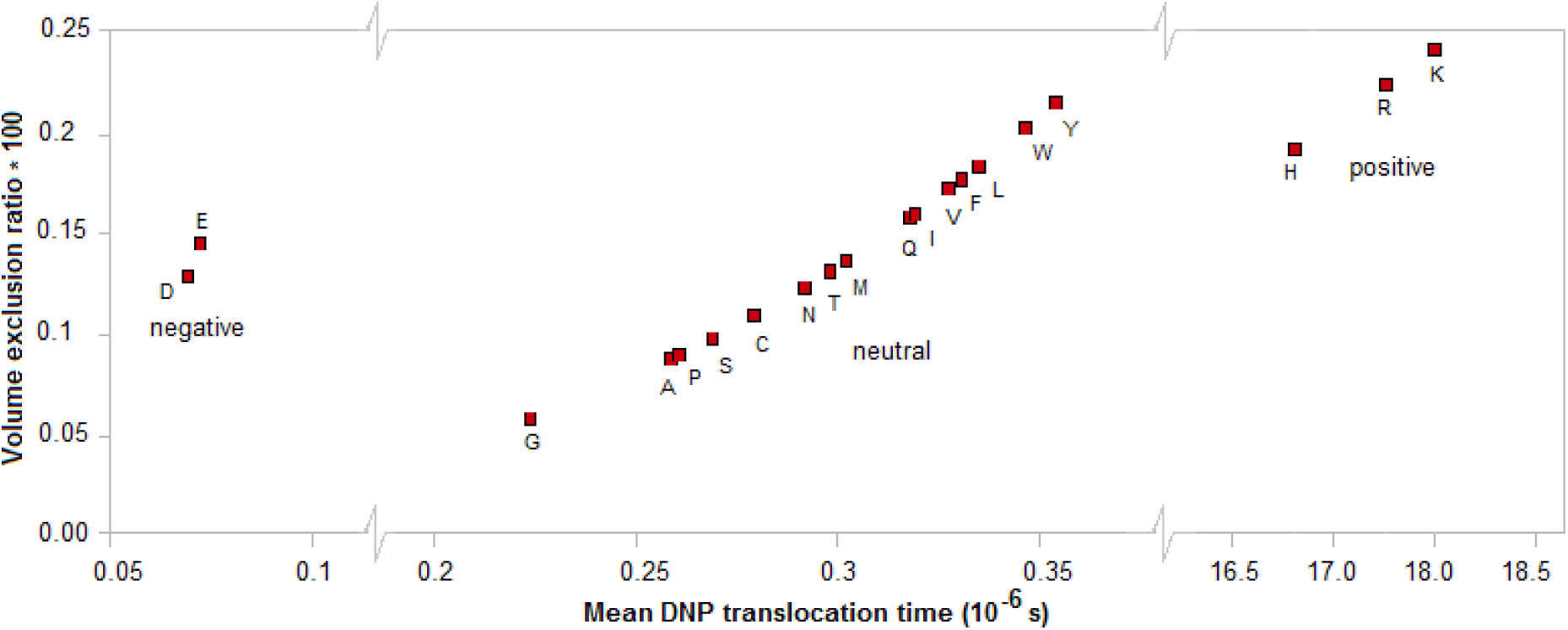
Scatter chart of mean of translocation time of particle in a tandem cell from time of entry into DNP (length L_34_ = 10 nm, negligible cross-section) to time of exit into *trans*2 vs volume exclusion ratio. V_05_ = 0.365 V, V_34_ = ~0.18 V.

**Figure 3.**
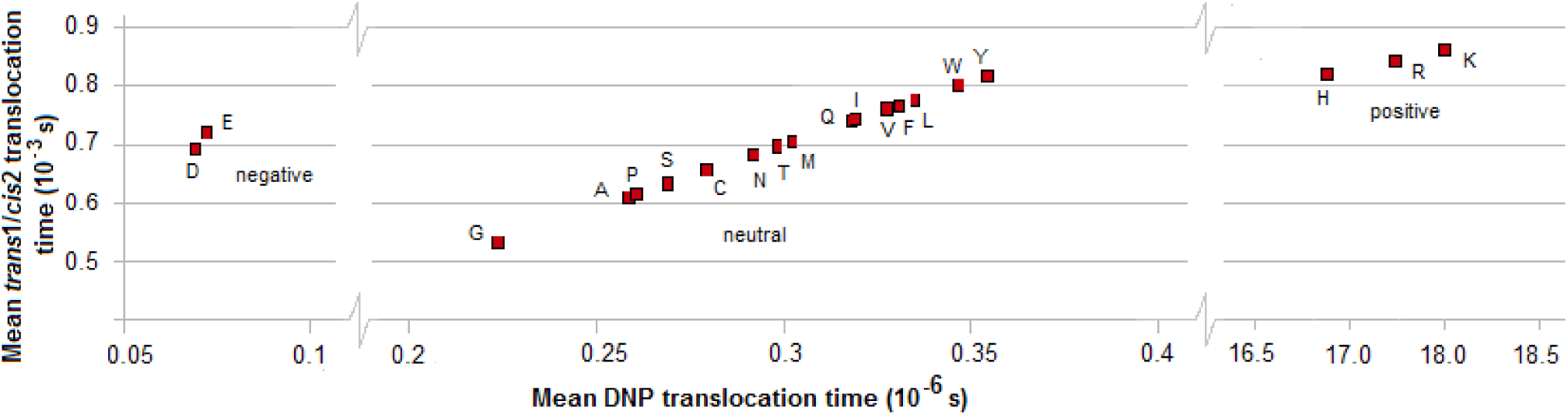
Scatter chart of mean of translocation time of particle from time of entry into DNP (length L_34_ = 10 nm, negligible cross-section) to time of exit into *trans*2 vs mean of time for particle to translocate from top of *trans*1/*cis*2 (length L_23_ = 1 μm, cross-section area = 1 μm^2^) to entrance of DNP. V_05_ = 0.365 V, V_23_ = ~1.6 mV, V_34_ = ~0.18 V.

The discriminators described above are computed measures. The experimentally measurable quantities are somewhat different. Thus,

1) Rather than T*_trans_*_1/*cis*2_ what is measured is the inter-arrival time between successive residues arriving at the pore. This quantity is T*_trans_*_1/*cis*2_ + T_gen_, where T_gen_ is the time for a residue to be generated at the top of *trans*1/*cis*2 and is, like T*_trans_*_1/*cis*2_, a random variable. In the tandem cell T_gen_ is replaced with the time T_c_ to cleave a residue from a peptide; see below. The inter-arrival time thus contains more information than T*_trans_*_1/*cis*2_ because the generation/cleaving time will vary with the amino acid;

2) The time spent by a residue inside DNP is more than the translocation time T_DNP_ because of the additional dwell time due to the reaction of the residue with the pore wall (which in a biological pore is a protein that may contain charged residues). Thus positively charged residues in the lumen will slow down negatively charged cleaved residues and vice versa, but neutral cleaved residues are not affected either way. Additional dwell time may result if a chemical adapter (similar to cyclodextrin in DNA sequencing [22]) is used to slow down the residue;

3) Current blockade, which reflects the change in the amplitude of the pore current from the baseline value, is, as mentioned above, determined by many more factors than volume exclusion. An important one is the presence of charged residues in the pore lumen (often by design; see, for example, [26]). The resulting electro-osmotic force may have a significant effect on the blockade level.

Wet experiments may be done with free amino acids in a tandem cell or single electrolytic cell to confirm (or not) the grouping and ordering seen in Figures 2 and 3.

### 5.3 Order of arrival of residue at DNP, occupancy in DNP, and minimum cleaving interval for exopeptidase

Two conditions need to be satisfied for accurate sequencing:

a. cleaved residues must enter DNP in natural order;
b. no more than one residue may occupy DNP at any time.

These conditions can be used to determine the minimum cleaving interval T_c_ between residues that are successively cleaved by the exopeptidase. Since cleaving behavior is stochastic and will vary with the amino acid, let T_c.min-X_ and T_c.max-X_ be the minimum and maximum cleaving times for amino acid X.

*Condition a*. Let residue X_1_ be cleaved at time t = 0. Its mean translocation time through *trans*1/*cis*2 is E(T*_trans_*_1/*cis*2-X1_) and standard deviation is σ*_trans_*_1/*cis*2-X1_. The next residue X_2_ is cleaved no earlier than t = T_c.min-X2_. Assuming 6σ support for the distribution, X_1_ arrives at the entrance to DNP latest by t = E(T*_trans_*_1/*cis*2-X1_) + 3σ*_trans_*_1/*cis*2-X1_. The earliest that X_2_ can arrive at DNP is t = T_c.min-X2_ + max (0, E(T*_trans_*_1/*cis*2-X2_) − 3σ*_trans_*_1/*cis*2-X2_). For X_2_ to follow X_1_ requires

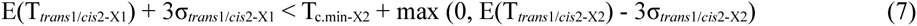

From the data in Table 1 in the Appendix, columns 8 and 9 (mean translocation time and standard deviation for *trans*1/*cis*2), max (0, E(T*_trans_*_1/*cis*2-X2_) − 3σ*_trans_*_1/*cis*2-X2_) = 0 for any amino acid. Equation 7 reduces to

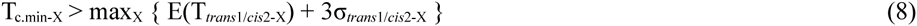

over all X. The maximum values occur for X = K (Lys), with E(T*_trans_*_1/*cis*2-X_) = 0.8632×10^−3^ and σ*_trans_*_1/*cis*2-X_ = 0.7077×10^−3^, leading to

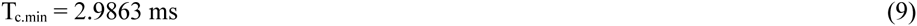

over all X.

*Condition b*. Consider residue X_1_ to be cleaved before X_2_. Since condition (a) has to be satisfied, X_1_ arrives at the entrance to DNP before X_2_. Let it arrive at time t = 0. The latest it can exit DNP is at time t = E(T_DNP-X1_) + 3σ_DNP-X1_. The earliest that X_2_ can arrive at the entrance of DNP is at t = T_c.min_ + max (0, E(T*_trans_*_1/*cis*2-X2_) − 3σ*_trans_*_1/*cis*2-X2_) = T_c.min_. Therefore for condition (b) to be satisfied

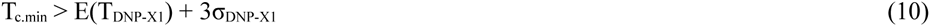

From Table 1 (columns 6 and 7), the maximum of the right hand side in Equation 10 occurs once again for X_1_ = K (Lys), with E(T_DNP-X1_) = 15.1215×10^−6^ and σ_DNP-X1_ = 15.0653×10^−6^, leading to

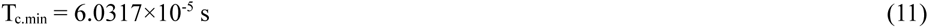

which is less than the value in Equation 9. Since Equation (9) has to be satisfied, it sets the minimum cleaving interval for any amino acid. (Thus Condition a subsumes Condition b.)

### 5.4 Behavior at an interface

The Fokker-Planck model mentioned above does not consider the behavior of the particle at the interface between two sections. In reality a particle oscillates at an interface because of diffusion. The effect of this on each type of residue, charged or not, is now considered.

a) *Negative residues* at the interface between *trans*1/*cis*2 and DNP experience a positive drift field inside both regions. Using formal probabilistic arguments [6] it can be shown that with sufficiently large V_05_ a negative residue will eventually pass into DNP, such passage being aided indirectly by the reflecting boundaries in *trans*1/*cis*2. (A cleaved residue cannot regress into UNP as the remaining peptide blocks its passage.) The behavior at the interface between DNP and *trans*2 is similar. The tapered geometry of *trans*1/*cis*2 shown in Figure 1 aids passage into DNP. Similarly the abrupt increase in cross-section from DNP to *trans*2 decreases the probability of a detected particle regressing into DNP from *trans*2.

b) *Positive residues* experience a negative drift field inside both regions. Because of this there is a non-zero probability that a positive residue may ultimately not enter DNP and therefore be ‘lost’ to diffusion in *trans*1/*cis*2. Or on entering DNP it may be trapped inside and neither regress into *trans*1/*cis*2 nor exit into *trans*2. One possible solution to the first problem is to design the pore lumen so as to prevent regression of the residue once it has entered DNP. One can also consider use of a hydraulic pressure gradient to prevent entry of a residue into DNP; however the hydrodynamic radius of an amino acid is too small for the pressure to be comparable to the electric field. (Compare this with the behavior of polyethylene glycol (PEG) in a nanopore with combined electric field and hydraulic pressure gradients [19]: 12 kDa PEG molecules with a length of 0.35 nm have a hydrodynamic radius of 3.2 nm, which is ~10 × average radius for an amino acid [24].) A third solution is to redo the sequencing using a second copy of the peptide with the voltage reversed (if the pore is ion-sensitive, one with the appropriate sense is to be used). In this case the roles of positive and negative residues are reversed. Thus positive residues are ‘lost’ to diffusion when V_05_ > 0 while negative residues are ‘lost’ to diffusion when V_05_ < 0. (In the latter case the header and trailer must have the appropriate charge sign.) With this approach two sequences are obtained with some or all positive residues missing in one and some or all negative residues missing in the other. Since the neutral residues are not affected the correct sequence can be obtained by merging the two individual sequences. However a residue that is trapped inside DNP and clogs it will still pose a problem. In this case there appears to be no alternative to re-sequencing with another copy of the peptide.

c) *Neutral residues* at the interface between *trans*1/*cis*2 and DNP are not affected by the electric field in either region. They are therefore subject entirely to diffusion. In this case the tapered geometry of *trans*1/*cis*2 in Figure 1 is useful in promoting entry from *trans*1/*cis*2 into DNP and also reduces the probability of permanent regression into *trans*1/*cis*2 from DNP. Although a hydraulic gradient could be used to assist entry into DNP, the improvement is minimal because its effect is small for reasonable values of hydraulic pressure, which are usually limited to 5-10 atm for solid-state membranes [19] (1 atm = 1.01325×10^5^ Pa).

The behavior at the interface between DNP and *trans*2 can be similarly understood, along with the fact that the abrupt change in diameter from DNP to *trans*2 acts as a deterrent to regression from *trans*2 into DNP.

### 5.5 Sample size requirements for reliable residue identification, confidence levels for a given sample size

The two time-based discriminators discussed above are mean values. To obtain a sample mean value which approaches the population (that is, calculated) mean for amino acid X, sequencing has to be done N (= sample size) times to distinguish the sample mean of X from that for another amino acid Z. The value of N, which depends on how close the mean translocation times of two amino acids are and the desired confidence level, can be calculated using standard formulas from statistics. Thus with a population mean E and standard deviation σ, margin of error e, and confidence level α (equivalently percentile value = 1 − α/2), the critical value Z_α/2_ of the normal distribution can be obtained from tables or calculated using statistical software (R was used in the present work). For example, with a confidence level of 0.95, α is 0.05, the percentile is 97.5, and the critical value is 1.96. The number of samples required for the sample mean E to approach the population mean within error e is

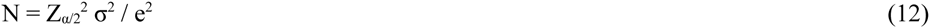

Tables 2 and 3 in the Appendix give the required sample sizes for DNP and *trans*1/*cis*2 for each amino acid X and its nearest neighbor (that is, the amino acid Z whose mean is closest to the mean of X) for three confidence levels: 90%, 80%, 70%. σ is taken from Table 1, e = k × min | E_X_ - E_Z_ | where Z is the amino acid in column 6 or 8 with mean E_Z_ nearest to the mean E_X_ for X, and k < 0.5. (This nearest neighbor can in almost all cases be identified visually in Figures 2 and 3, where the amino acids separate into ordered groups.) Figures 4 and 5 show histograms of the sample size for DNP and *trans*1/*cis*2 respectively for k = 0.4.

**Figure 4.**
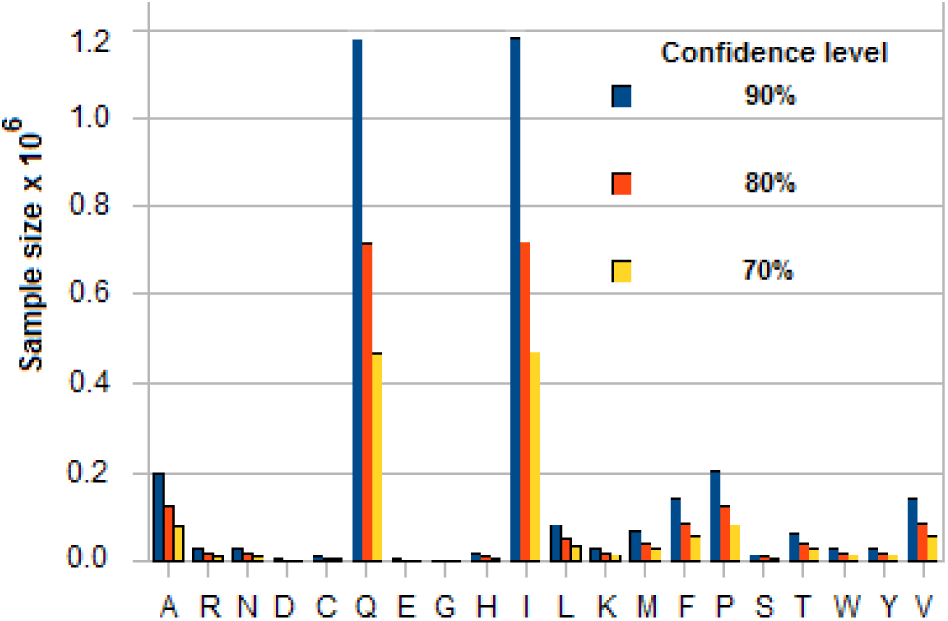
Sample sizes for three confidence levels using standard statistical formula for each amino acid based on standard deviation of its translocation time in DNP and the margin of error (= 0.4 × smallest difference between the amino acid’s mean translocation time and that of any of the other 19). See Tables 1 and 2 in Appendix for calculated data.

**Figure 5.**
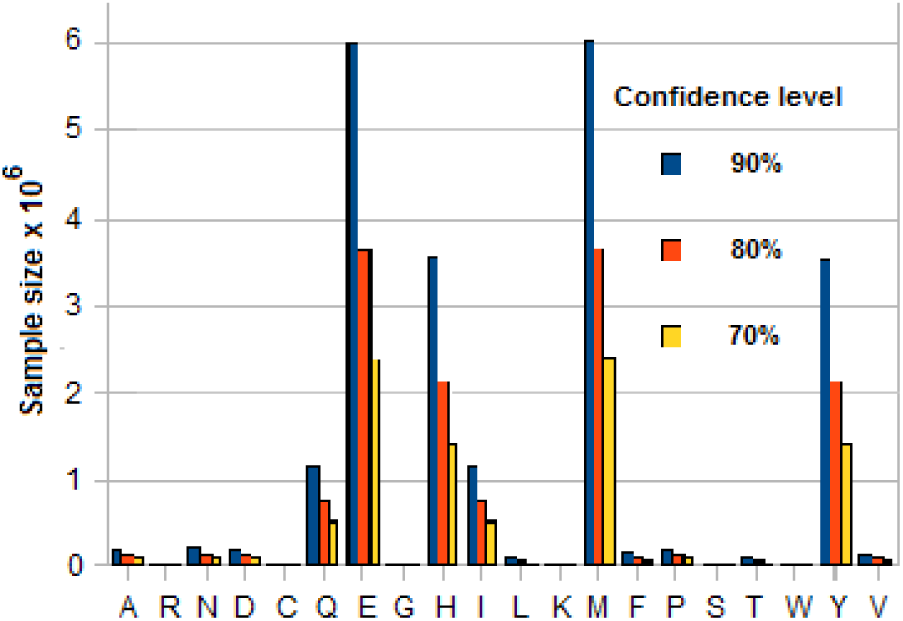
Sample sizes for three confidence levels using standard statistical formula for each amino acid based on standard deviation of its translocation time in *trans*1/*cis*2 and the margin of error (= 0.4 × smallest difference between the amino acid’s mean translocation time and that of any of the other 19). See Tables 1 and 3 in Appendix for calculated data.

The value of N to use in the sequencing is the largest sample size N_max_ over all the amino acids. In determining N_max_ the discriminator to use for an amino acid is based on the smallest number of samples over all its discriminators. For example, Asn (symbol N) has E(T_DNP_) = ~0.273× 10^−6^ which is 0.0055 ×10^−6^ from the mean time of Thr (T) and requires ~27800 samples for a confidence level of 90%. It has E(T*_trans_*_1/*cis*2_) = 0.683× 10^−3^ which is 0.005× 10^3^ from the mean time of Asp (D) and requires > 200000 samples. The discriminator to use for Asn is therefore E(T_DNP_).

Amino acid pairs whose mean times are very close to each other are the ones that effectively determine N_max_. As seen from Tables 2 and 3 (or Figures 2 and 3) the problem pairs are Ala (A) - Pro (P), Gln (Q) - Ile (I), and Phe (F) - Val (V), all with N values close to 10^6^ (DNP) or far in excess of it (*trans*1/*cis*2). A more manageable value of N_max_ is possible if these highly error-prone residue pairs are excluded from its determination. This lowers the confidence levels for their measured means but their identification can be handled through error correction (which is made easier by the ordering property; see Figures 2 and 3; error correction could also be based on, for example, methods used in mass spectrometry [7]). This leads to N_max_ = ~81000 for a confidence level of 90% or better for the other 14 amino acids, and ~32000 for a confidence level of 70% or better. For a long peptide, N_max_ could in principle be lowered by a factor of L_pep_/20, where L_pep_ is the length, because of repeats; this assumes that the 20 amino acids occur in proteins with equal probability.

Conversely for a given maximum number of samples N_max_ one can find the confidence level for the sample mean of an amino acid X to be no farther from the population mean than e = k × min | E_X_ − E_Z_ |, where e is the distance to the nearest mean, with k < 0.5. This can be obtained from the critical value using the statistical formula

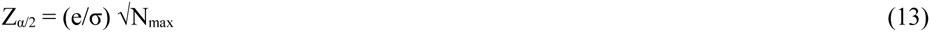

and tables (or the *pnorm* function in the software package R). For example, with DNP and N = 10000 consider X = A (Ala) with σ = 0.199021× 10^−6^. Its nearest mean neighbor Z = P (Pro) with distance to mean of Z = 0.001833× 10^−6^. With k = 0.4 the resulting critical value Z_α/2_ = 0.3684, for which the confidence level is 0.3976 (39.76%). Table 4 in the Appendix gives the confidence levels for the 20 amino acids for k = 0.4 and N = 10000 in DNP and *trans*1/*cis*2. Figure 6 shows a histogram of comparative confidence levels of residue identification in DNP and *trans*1/*cis*2 for all 20 amino acids.

**Figure 6.**
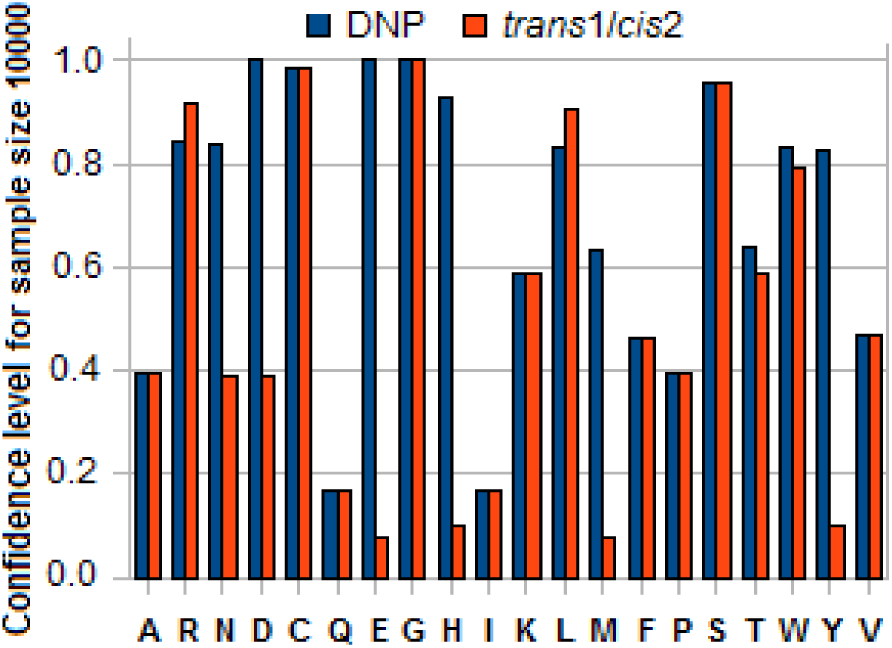
Histogram of confidence levels for an amino acid X to ensure that sample mean translocation time in DNP or *trans*1/*cis*2 is within 0.4 × smallest difference between the calculated mean for X and that for any of the other 19 for a sample size of 10000. See Table 4 in Appendix.

Assuming ergodicity and the availability of a sufficient quantity of the assay sample a parallel implementation of N_max_ tandem cells can be used with N_max_ copies of the peptide to quickly obtain the sample mean for every residue in the peptide. Such an approach (which will have to be automated because of the large values of N_max_ involved) would be more appropriate to research than to clinical or forensic assays where only a limited amount of the test sample may be available. Alternatively N× sequencing with one copy of the peptide may be possible by recycling the cleaved residues after their detection in the tandem cell back into *cis*1 for translocation through UNP and *trans*1/*cis*2 to DNP for another round of detection. This recycling can be done N_max_ times; it assumes that the recycled residues are not affected by the exopeptidase attached to UNP. For short peptides the value of N_max_ can be set adaptively after the first few sample runs have yielded a tentative sequence. A tandem cell with recycling capability that uses a hydraulic gradient to ‘pump’ detected residues back to *cis*1 is currently being designed, details will be available later.

By way of comparison, the reduced value of _Nmax_ (~80000) is of the same order as the number of crystals used in serial femtosecond nanocrystallography (SFX) [27] to determine protein structure using a ‘diffract-then-destroy’ approach. SFX uses a high intensity laser pulse to capture the diffraction image of one of ~10^4^ crystals in a liquid jet (LJ-SFX) injected by a nozzle into the path of the laser or a fixed target (FT-SJX) interposed mechanically. The entire sample is destroyed in the process, but not before the image is captured.

## 6 Discussion

The feasibility of the proposed scheme depends crucially on the exopeptidase being able to cleave every residue in the peptide in a reasonable amount of time (Equation 9 gives the minimum cleaving interval for typical parameter values). Assuming that this condition is satisfied, the following factors may be considered in a practical implementation:

1. With translocation times through DNP on the order of 10^−7^ s (see Table 1), the bandwidth required is ~10 MHz (including noise filtering). The lower signal-to-noise ratio in this frequency range combined with the pA-level blockade levels and fast translocation makes detection difficult (this is a problem in nanopore-based sequencing of any analyte, including DNA). Methods to slow down translocation in a tandem cell are discussed in [6]. Another method described in the literature uses charged residues in the pore lumen to slow down a translocating charged analyte [26]. Thus positively charged residues in the lumen will slow down negatively charged analyte residues and vice versa, but have no effect on neutral analytes.
2. As noted earlier a variety of methods based on Hidden Markov Models, Viterbi algorithms, and complex neural nets [17,18] have been used to increase base calling accuracy in strand sequencing of DNA. These methods, which are designed to work with the current signal record, can be modified to work with multiple discriminators (Section 5.1) to improve residue calling accuracy.
3. The proposed scheme assumes that with amino (carboxyl) exopeptidase the peptide enters UNP N-terminal (C-terminal) first. There is no guarantee that this will happen. Entry of the wrong end can be detected when cleaving fails to occur, as recognized by the absence of the characteristic blockades that would occur due to cleaved residues. In this case the intact peptide entering *trans*2 can be recycled to *cis*1 for another attempt at detection, to be repeated until residue-driven blockades are detected. With two identical copies of the peptide, two sequencers, one with amino exopeptidase and the other with carboxyl, can be used to increase the probability of successful sequencing. An alternative approach that dispenses with any dependence on the peptide’s random orientation when entering DNP may be based on two tandem cells in tandem, the first with amino exopeptidase and the second with carboxyl. The device would then have the structure [*cis*1, UNP with amino peptidase, *trans*1/*cis*2, DNP with carboxyl peptidase, *trans*2/*cis*3, third (sensing) nanopore (TNP), *trans*3]. To guarantee detection in the second stage of a peptide that was not sequenced in DNP because it entered UNP C-terminal first, the unsequenced polymer has to enter DNP C-terminal first. This can be ensured if the poly-X leader (which entered UNP C-terminal first) is longer than the length of *trans*1/*cis*2 so that the trailing polymer is still inside UNP and the leader (with its free C-terminal in front) enters DNP C-terminal first. (High enough voltages that are within the breakdown limit may ensure such entry. Up to 0.7 V can be applied across a biological nanopore of length 10 nm [6].) This ensures that the leading residue is cleaved by the carboxyl exopeptidase attached to the downstream side of DNP. When sequencing occurs in the first stage spurious signals from cleaved residues that try to enter TNP after detection in DNP can be avoided by flushing them out after they have entered *trans*2 (thus effectively deactivating TNP). Yet another possible, and somewhat simpler, alternative (although it requires an additional step) is to attach a capping molecule (similar to a biotin-streptavidin tether [28]) to the trailer at either the C-end or N-end of the peptide to prevent that end from entering UNP.
4. A folded protein could be loaded into the tandem cell and unfolded by an enzyme (unfoldase) like ClpX [9] before sequencing. The unfoldase, which acts as a motor that both unfolds and translocates the protein, could be attached to the upstream side of UNP in *cis*1 so that the protein enters UNP unfolded (and is then cleaved by the exopeptidase attached to the downstream side of UNP). Alternatively the unfoldase could be attached to the downstream side (similar to [9]) of a precursor nanopore in a double tandem cell with the structure [*cis*0, precursor UNP with ClpX, *trans*0/*cis*1, UNP with exopeptidase, *trans*1/*cis*2, DNP, *trans*2]. In this case the unfolded protein translocates to UNP after it has passed through ClpX, following which its behavior would be similar to that in the basic tandem cell. In this unfold-cleave-sequence approach, the reaction rates of the unfolding and cleaving enzymes have to be matched (balanced) to prevent stalling or clogging in UNP.
5. The optimum peptide length handled by an efficient mass spectrometer is ~20 [7]. Considerably larger lengths may be possible with a tandem cell if a practical version can be designed to match the performance of the theoretical model. If in addition unfolding can be implemented as in (4) above, the tandem cell could be used to construct the primary sequence of a whole protein.
6. If an amino acid can be uniquely identified by a transverse recognition tunneling (RT) current as in [8], a cascade of 21 nanopores may be used to fully sequence a peptide. In such a tandem cascade the first tandem stage is used to cleave residues in the peptide, followed by 20 pores each of which is designed to recognize a unique amino acid. Such a system can sequence a peptide without having to depend on ionic current blockades and the extreme measurement precision required to distinguish among their closely spaced values in the presence of noise. Alternatively a single DNP with 20 recognizers and 20 pairs of transverse electrodes may also be possible. In either case the length of the pore is no longer a crucial issue as it is in most nanopore sequencing approaches to date. Correlations among the 20 transverse current records can be used not only to improve residue calling accuracy but also to extract other kinds of peptide-related information. The order of the recognizers may also be optimized to maximize discrimination among the residues.
7. It is possible for some neutral residues to attract ions in an electrolyte and carry a resulting charge [19,29]. A cleaved residue that is ordinarily neutral can therefore become positively or negatively charged due to formation of an anion or cation complex. No information is available about whether amino acids form such complexes in aqueous KCl or not, so this line of investigation has not been pursued.
8. The tandem cell approach to peptide sequencing as described above is a destructive process as the peptide is broken down into its constituent amino acids. Unlike exonuclease-based DNA sequencing, where re-sequencing of the original strand from the cleaved bases can be done using the individual cleaved nucleotides and a template with an enzyme motor attached to a nanopore [16], there is no simple way to re-synthesize the peptide that can be integrated with the tandem cell. However as noted toward the end of Section 5.5, by routing cleaved residues that have translocated into *trans*2 after detection in DNP into the same or a second tandem cell they can be sequenced N_max_ times in a loop. Such re-sequencing can also be viewed as N× coverage (something that is normally done for error checking, especially in genome sequencing).
9. For other implementation-related issues in sequencing with a tandem cell, such as voltage drift and monomers that might stick to channel walls, and their possible resolution, see discussion in [6].

### A note on two-pore systems in protein analysis

There appears to be one other reported instance in the literature of a system with twin nanopores for protein analysis. In [30] two nanopores in series are used to measure mobility and particle sizes to identify specific proteins. The nanopores are comparatively larger, with cross-section dimensions that are several 10’s of nm. The system is structurally and procedurally different from the tandem cell described here. (For two-pore systems used in DNA sequencing see Supplement to [6].)

## Appendix

**Table 1.** Statistics of amino acid translocation times in DNP and *trans*1/*cis*2

| Amino acid | Abbrevn and Charge<sup>(a)</sup> | Hydrodynamic radius<sup>(b)</sup><br>R<sub>aa</sub> (10<sup>-10</sup> m) | Diffusion coefficient<sup>(c)</sup><br>D<sub>aa</sub> (10<sup>-10</sup> m<sup>2</sup>/s) | Mobility<sup>(d)</sup><br>μ<sub>aa</sub> (10<sup>-8</sup> m/Vs) | Mean translocation time in DNP<sup>(e)</sup><br>(10<sup>-6</sup> s) | Std deviation of translocation time in DNP<sup>(f)</sup><br>(10<sup>-6</sup> s) | Mean translocation time in *trans1/cis2*<sup>(g)</sup><br>(10<sup>-3</sup> s) | Std deviation of translocation time in *trans1/cis2*<sup>(h)</sup><br>(10<sup>-3</sup> s) | Volume exclusion ratio<sup>(i)</sup><br>(× 100) |
| --- | --- | --- | --- | --- | --- | --- | --- | --- | --- |
| Ala | A n | 2.6600 | 8.2052 | 3.1950 | 0.2437490 | 0.199021 | 0.6093730 | 0.4975510 | 0.086291 |
| Arg | R + | 3.6000 | 6.0627 | 2.3607 | 14.7526920 | 14.697848 | 0.8421130 | 0.6904320 | 0.219551 |
| Asn | N n | 2.9800 | 7.3241 | 2.8519 | 0.2730730 | 0.222963 | 0.6826810 | 0.5574070 | 0.122299 |
| Asp | D - | 3.0200 | 7.2271 | 2.8141 | 0.0677110 | 0.033888 | 0.6776980 | 0.5510350 | 0.127425 |
| Cys | C n | 2.8600 | 7.6314 | 2.9715 | 0.2620760 | 0.213984 | 0.6551910 | 0.5349610 | 0.107778 |
| Gln | Q n | 3.2300 | 6.7572 | 2.6311 | 0.2959810 | 0.241668 | 0.7399530 | 0.6041690 | 0.156806 |
| Glu | E - | 3.1400 | 6.9509 | 2.7066 | 0.0704010 | 0.035234 | 0.7046270 | 0.5729310 | 0.143695 |
| Gly | G n | 2.3200 | 9.4076 | 3.6632 | 0.2125930 | 0.173582 | 0.5314840 | 0.4339540 | 0.056826 |
| His | H + | 3.4900 | 6.2538 | 2.4351 | 14.3019160 | 14.248747 | 0.8163820 | 0.6693360 | 0.199341 |
| Ile | I n | 3.2400 | 6.7363 | 2.6230 | 0.2968980 | 0.242416 | 0.7422440 | 0.6060400 | 0.158312 |
| Leu | L n | 3.3900 | 6.4383 | 2.5070 | 0.3106430 | 0.253639 | 0.7766070 | 0.6340970 | 0.182135 |
| Lys | K + | 3.6900 | 5.9148 | 2.3031 | 15.1215100 | 15.065294 | 0.8631660 | 0.7076930 | 0.237124 |
| Met | M n | 3.0800 | 7.0863 | 2.7593 | 0.2822360 | 0.230445 | 0.7055900 | 0.5761120 | 0.135391 |
| Phe | F n | 3.3500 | 6.5151 | 2.5369 | 0.3069780 | 0.250646 | 0.7674440 | 0.6266150 | 0.175553 |
| Pro | P n | 2.6800 | 8.1439 | 3.1711 | 0.2455820 | 0.200517 | 0.6139550 | 0.5012920 | 0.088293 |
| Ser | S n | 2.7600 | 7.9079 | 3.0792 | 0.2529130 | 0.206502 | 0.6322820 | 0.5162560 | 0.096624 |
| Thr | T n | 3.0400 | 7.1795 | 2.7956 | 0.2785710 | 0.227452 | 0.6964270 | 0.5686300 | 0.130043 |
| Trp | W n | 3.5000 | 6.2359 | 2.4282 | 0.3207230 | 0.261869 | 0.8018070 | 0.6546730 | 0.201122 |
| Tyr | Y n | 3.5700 | 6.1136 | 2.3806 | 0.3271370 | 0.267106 | 0.8178430 | 0.6677660 | 0.213903 |
| Val | V n | 3.3200 | 6.5740 | 2.5598 | 0.3042290 | 0.248402 | 0.7605710 | 0.6210040 | 0.170728 |

(a) n = neutral (b) Values from [24] (c), (d) Values computed from Equation 6 in main text (e), (f), (g), (h) Values computed from Equations 1-4 in main text (i) Values computed from Equation 5 in main text; L = L<sub>34</sub> = 10 nm, r = 1.5 nm

**Table 2.** Histogram of amino acid sample sizes for three confidence levels (DNP)

| DNP |  | Confidence level = 0.9 |  | Confidence level = 0.8 |  | Confidence level = 0.7 |  |
| --- | --- | --- | --- | --- | --- | --- | --- |
| Amino Acid | Nearest Amino Acid<sup>(a)</sup> | Difference in means (10<sup>-6</sup> s) | Sample size | Difference in means (10<sup>-6</sup> s) | Sample size | Difference in means (10<sup>-6</sup> s) | Sample size |
| A | P | 0.001833 | 199346 | 0.001833 | 121011 | 0.001833 | 79147 |
| R | K | 0.368818 | 26854 | 0.368818 | 16301 | 0.368818 | 10662 |
| N | T | 0.005498 | 27809 | 0.005498 | 16881 | 0.005498 | 11041 |
| D | E | 0.002690 | 2683 | 0.002690 | 1629 | 0.002690 | 1065 |
| C | S | 0.009163 | 9221 | 0.009163 | 5598 | 0.009163 | 3661 |
| Q | I | 0.000917 | 1174449 | 0.000917 | 712938 | 0.000917 | 466296 |
| E | D | 0.002690 | 2901 | 0.002690 | 1761 | 0.002690 | 1151 |
| G | A | 0.031156 | 524 | 0.031156 | 318 | 0.031156 | 208 |
| H | R | 0.450776 | 16895 | 0.450776 | 10256 | 0.450776 | 6708 |
| I | Q | 0.000917 | 1181730 | 0.000917 | 717358 | 0.000917 | 469187 |
| L | F | 0.003665 | 80987 | 0.003665 | 49162 | 0.003665 | 32154 |
| K | R | 0.368818 | 28214 | 0.368818 | 17127 | 0.368818 | 11201 |
| M | T | 0.003665 | 66853 | 0.003665 | 40582 | 0.003665 | 26542 |
| F | V | 0.002749 | 140574 | 0.002749 | 85334 | 0.002749 | 55812 |
| P | A | 0.001833 | 202354 | 0.001833 | 122837 | 0.001833 | 80341 |
| S | P | 0.007331 | 13417 | 0.007331 | 8144 | 0.007331 | 5327 |
| T | M | 0.003665 | 65127 | 0.003665 | 39535 | 0.003665 | 25857 |
| W | Y | 0.006414 | 28186 | 0.006414 | 17110 | 0.006414 | 11191 |
| Y | W | 0.006414 | 29325 | 0.006414 | 17801 | 0.006414 | 11643 |
| V | F | 0.002749 | 138068 | 0.002749 | 83813 | 0.002749 | 54817 |

<sup>(a)</sup> Amino acid with closest mean translocation time

**Table 3.** Histogram of amino acid sample sizes for three confidence levels (*trans*1/*cis*2)

| *trans1/cis2* |  | Confidence level = 0.9 |  | Confidence level = 0.8 |  | Confidence level = 0.7 |  |
| --- | --- | --- | --- | --- | --- | --- | --- |
| Amino Acid | Nearest Amino Acid <sup>(a)</sup> | Difference in means ($10^{-3}$ s) | Sample size | Difference in means ($10^{-3}$ s) | Sample size | Difference in means ($10^{-3}$ s) | Sample size |
| A | P | 0.004582 | 199388 | 0.004582 | 121036 | 0.004582 | 79163 |
| R | K | 0.021053 | 18186 | 0.021053 | 11039 | 0.021053 | 7220 |
| N | D | 0.004983 | 211591 | 0.004983 | 128444 | 0.004983 | 84008 |
| D | N | 0.004983 | 206781 | 0.004983 | 125524 | 0.004983 | 82099 |
| C | D | 0.022507 | 9553 | 0.022507 | 5799 | 0.022507 | 3792 |
| Q | I | 0.002291 | 1175984 | 0.002291 | 713870 | 0.002291 | 466905 |
| E | M | 0.000963 | 5985312 | 0.000963 | 3633327 | 0.000963 | 2376372 |
| G | A | 0.077889 | 524 | 0.077889 | 318 | 0.077889 | 208 |
| H | Y | 0.001461 | 3549136 | 0.001461 | 2154469 | 0.001461 | 1409127 |
| I | Q | 0.002291 | 1183278 | 0.002291 | 718298 | 0.002291 | 469802 |
| L | F | 0.009163 | 80978 | 0.009163 | 49157 | 0.009163 | 32151 |
| K | R | 0.021053 | 19107 | 0.021053 | 11598 | 0.021053 | 7586 |
| M | E | 0.000963 | 6051959 | 0.000963 | 3673785 | 0.000963 | 2402833 |
| F | V | 0.006873 | 140554 | 0.006873 | 85322 | 0.006873 | 55804 |
| P | A | 0.004582 | 202397 | 0.004582 | 122863 | 0.004582 | 80358 |
| S | P | 0.018327 | 13417 | 0.018327 | 8145 | 0.018327 | 5327 |
| T | E | 0.008200 | 81314 | 0.008200 | 49361 | 0.008200 | 32284 |
| W | H | 0.014575 | 34116 | 0.014575 | 20710 | 0.014575 | 13545 |
| Y | H | 0.001461 | 3532505 | 0.001461 | 2144374 | 0.001461 | 1402525 |
| V | F | 0.006873 | 138048 | 0.006873 | 83800 | 0.006873 | 54809 |

<sup>(a)</sup> Amino acid with closest mean translocation time

**Table 4.** Confidence level for sample mean of an amino acid to be within specified error for a given sample size

| Amino Acid | DNP |  |  |  | *trans1/cis2* |  |  |  |
| --- | --- | --- | --- | --- | --- | --- | --- | --- |
| | Nearest Amino Acid <sup>(a)</sup> | Difference in means ($10^{-6}$ s) | $Z_{a/2}$ <sup>(b)</sup> | Confidence level | Nearest Amino Acid <sup>(a)</sup> | Difference in means ($10^{-3}$ s) | $Z_{a/2}$ <sup>(b)</sup> | Confidence level |
| A | P | 0.001833 | 0.368403 | 0.397634 | P | 0.004582 | 0.368364 | 0.397595 |
| R | K | 0.368818 | 1.003733 | 0.844245 | K | 0.021053 | 1.219700 | 0.915457 |
| N | T | 0.005498 | 0.986352 | 0.836958 | D | 0.004983 | 0.357584 | 0.386933 |
| D | E | 0.002690 | 3.175165 | 0.999993 | N | 0.004983 | 0.361719 | 0.391033 |
| C | S | 0.009163 | 1.712838 | 0.984578 | D | 0.022507 | 1.682889 | 0.982686 |
| Q | I | 0.000917 | 0.151778 | 0.169957 | I | 0.002291 | 0.151679 | 0.169848 |
| E | D | 0.002690 | 3.053868 | 0.999984 | M | 0.000963 | 0.067233 | 0.075750 |
| G | A | 0.031156 | 7.179546 | 1.000000 | A | 0.077889 | 7.179471 | 1.000000 |
| H | R | 0.450776 | 1.265447 | 0.926484 | Y | 0.001461 | 0.087310 | 0.098269 |
| I | Q | 0.000917 | 0.151310 | 0.169441 | Q | 0.002291 | 0.151211 | 0.169332 |
| L | F | 0.003665 | 0.577987 | 0.586298 | F | 0.009163 | 0.578019 | 0.586324 |
| K | R | 0.368818 | 0.979252 | 0.833908 | R | 0.021053 | 1.189951 | 0.907595 |
| M | T | 0.003665 | 0.636160 | 0.631702 | E | 0.000963 | 0.066862 | 0.075333 |
| F | V | 0.002749 | 0.438706 | 0.465021 | V | 0.006873 | 0.438738 | 0.465051 |
| P | A | 0.001833 | 0.365655 | 0.394924 | A | 0.004582 | 0.365615 | 0.394884 |
| S | P | 0.007331 | 1.420035 | 0.955381 | P | 0.018327 | 1.419993 | 0.955375 |
| T | M | 0.003665 | 0.644532 | 0.637971 | E | 0.008200 | 0.576825 | 0.585359 |
| W | Y | 0.006414 | 0.979727 | 0.834114 | H | 0.014575 | 0.890521 | 0.792109 |
| Y | W | 0.006414 | 0.960518 | 0.825656 | H | 0.001461 | 0.087516 | 0.098500 |
| V | F | 0.002749 | 0.442670 | 0.468705 | F | 0.006873 | 0.442702 | 0.468734 |

<sup>(a)</sup> Amino acid with closest mean translocation time
<sup>(b)</sup> Critical value of normal distribution

### For comparison: Discriminator data for nucleotides

**Table 5.**
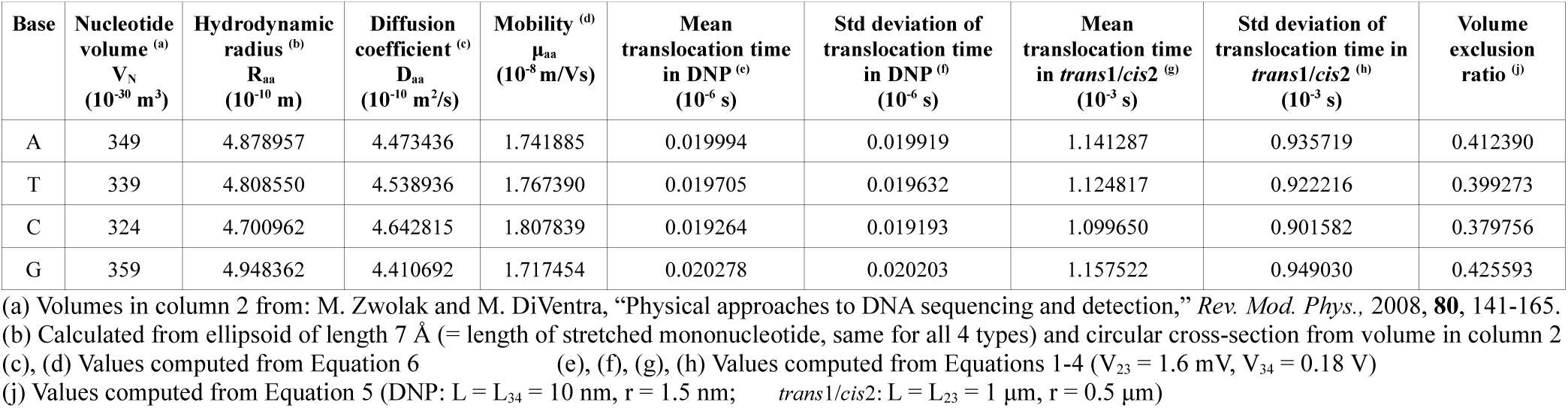
Statistics of translocation times in DNP and *trans*1/*cis*2 for the four nucleotide types

Figure 7 Combined scatter charts of translocation time through DNP vs translocation time through *trans*1/*cis*2 and of translocation time through DNP vs volume exclusion ratio

**Figure 7.**
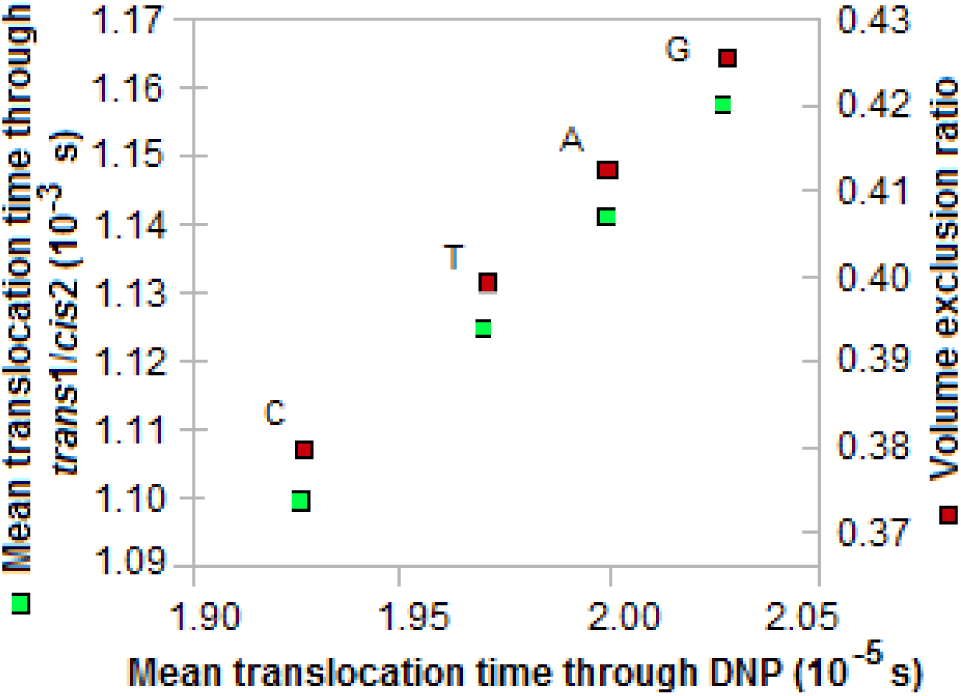
Scatter charts for nucleobases

Figure 8a Histogram of sample sizes for three confidence levels (DNP)

**Figure 8.**
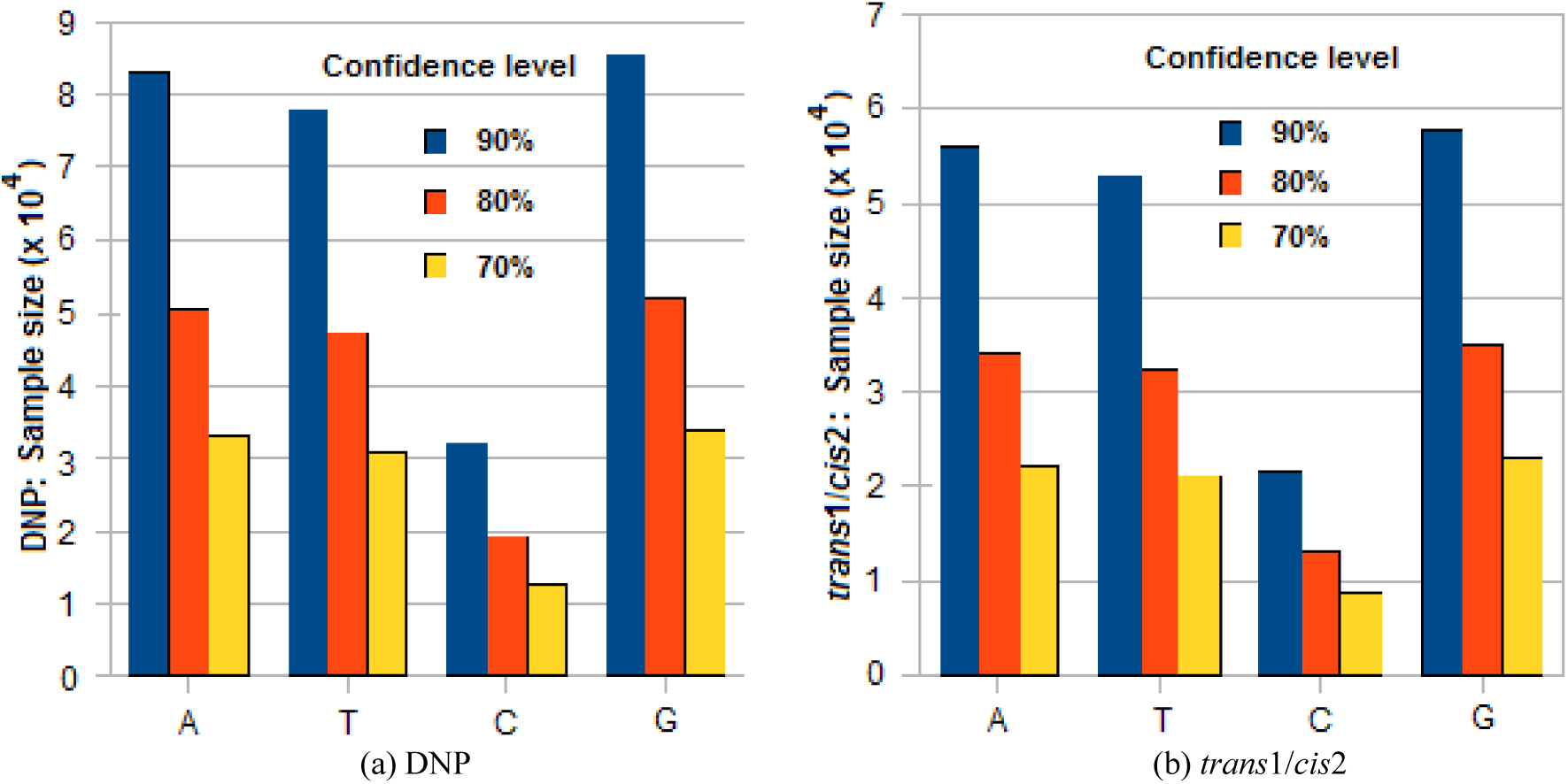
Histograms of sample sizes for three confidence levels

Figure 8b Histogram of sample sizes for three confidence levels (*trans*1/*cis*2)

Figure 9 Combined histograms of confidence levels for DNP and *trans*1/*cis*2 for sample size 10000

**Figure 9.**
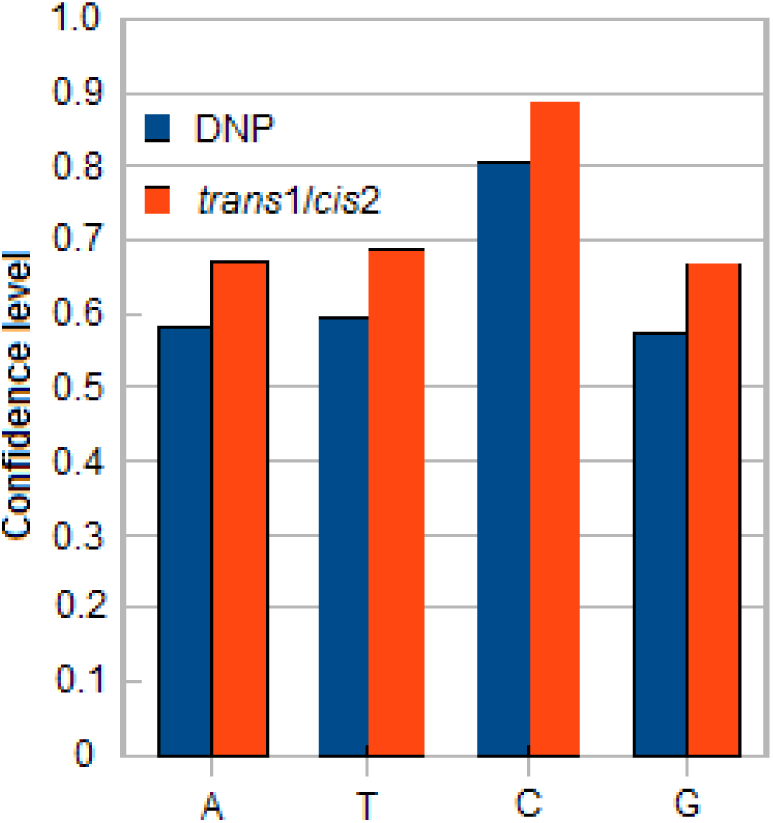
Histograms of confidence levels for sample size = 10000

